# Localization of BCR-ABL to stress granules contributes to its oncogenic function

**DOI:** 10.1101/801555

**Authors:** Sayaka Kashiwagi, Yoichiro Fujioka, Takeshi Kondo, Aya O. Satoh, Aiko Yoshida, Mari Fujioka, Hitoshi Sasajima, Maho Amano, Takanori Teshima, Yusuke Ohba

## Abstract

The oncogenic tyrosine kinase BCR-ABL activates a variety of signaling pathways and plays a causative role in the pathogenesis of chronic myelogenous leukemia (CML); however, the subcellular distribution of this chimeric protein remains controversial. Here, we report that BCR-ABL is localized to stress granules and that its granular localization contributes to BCR-ABL-dependent leukemogenesis. BCR-ABL-positive granules were not colocalized with any markers for membrane-bound organelles but were colocalized with HSP90a, a component of RNA granules. The number of such granules increased with thapsigargin treatment, confirming that the granules were stress granules. Given that treatment with the ABL kinase inhibitor imatinib and elimination of the N-terminal region of BCR-ABL abolished granule formation, kinase activity and the coiled-coil domain are required for granule formation. Whereas wild-type BCR-ABL rescued the growth defect in IL-3-depleted Ba/F3 cells, mutant BCR-ABL lacking the N-terminal region failed to do so. Moreover, forced tetramerization of the N-terminus-deleted mutant could not restore the growth defect, indicating that granule formation, but not tetramerization, through its N-terminus is critical for BCR-ABL-dependent oncogenicity. Our findings together provide new insights into the pathogenesis of CML by BCR-ABL and open a window for developing novel therapeutic strategies for this disease.

## INTRODUCTION

Chronic myelogenous leukemia (CML) is a myeloproliferative disorder resulting from a reciprocal translocation between chromosomes 9 and 22, which forms the characteristic Philadelphia (Ph^1^) chromosome. This translocation fuses the breakpoint cluster region (*BCR*) gene into the juxtaposition of the nonreceptor tyrosine kinase-encoding *ABL* gene to generate the fusion gene *BCR-ABL*, a driver mutation in CML. The fusion protein BCR-ABL, a constitutively activated tyrosine kinase, thereby plays a causative role in the pathogenesis of CML through the upregulation of a wide range of signaling pathways. It has been shown that expression of BCR-ABL can transform mouse fibroblast cells, factor-dependent hematopoietic cell lines, and primary bone marrow cells (Ren, 2005). Furthermore, the forced expression of BCR-ABL in mouse bone marrow cells results in a CML-like myeloproliferative disorder (Daley *et al.*, 1990). Such transformation activity of BCR-ABL is completely dependent on its tyrosine kinase activity. In fact, oncogenesis by BCR-ABL was diminished by the introduction of a point mutation in the ATP-binding site of ABL, which inactivates its kinase activity (Zhang and Ren, 1998). Based on these findings, tyrosine kinase inhibitors of ABL have been developed and utilized for CML treatment. For instance, the first generation of the tyrosine kinase inhibitor imatinib has innovated the treatment of CML and changed this disease into being controllable by oral administration (Druker *et al.*, 1996; Druker *et al.*, 2002).

The ABL kinase activity of BCR-ABL is necessary, as described above, but not sufficient for CML pathogenesis. Fusion with the BCR protein provides ABL kinase domains and motifs that are required for constitutive activity, interaction with other downstream signaling pathways, and subsequent oncogenic activities. For example, it was shown that mutant c-ABL activated by SH3 deletion cannot induce myeloproliferative disorder in mice (Gross *et al.*, 1999). A mutant p210 BCR-ABL lacking the N-terminal coiled-coil (CC) oligomerization domain is unable to induce myeloproliferative disorder in mice (He *et al.*, 2002).

Phosphorylation of tyrosine at 177 in BCR is also essential for the activation of Ras/extracellular-regulated kinase (ERK) and phosphoinositide 3-kinase (PI3K)/Akt pathways through binding to the SH2 domain of an adaptor protein, growth factor receptor-bound protein 2 (Grb2) (Sattler *et al.*, 2002). However, knowledge about the importance of other domains of BCR-ABL in oncogenic signaling pathways has been limited (Ren, 2005).

The spatiotemporal regulation of signaling molecules is needed to determine the diversity and specificity of intracellular signaling. Although BCR-ABL is predominantly localized in the cytoplasm, the specific subcellular localization of this fusion has also been reported. For example, Wertheim and colleagues demonstrated that BCR-ABL is localized to F-actin through the actin-binding domain in its C-terminus (Wertheim *et al.*, 2003). Another group also claimed that BCR-ABL was localized in the Golgi apparatus (Miroshnychenko *et al.*, 2010). In addition, both endogenous and exogenous BCR-ABL form granule-like structures in either hematopoietic or nonhematopoietic cells (Skourides *et al.*, 1999; Patel *et al.*, 2008).

Recent developments in imaging techniques have enabled us to examine the precise spatiotemporal regulation of intracellular signaling in living cells. We previously developed the Förster resonance energy transfer (FRET)-based biosensor Pickles (phosphorylation indicator of CrkL *en* substrate) and succeeded in measuring BCR-ABL kinase activity and its drug response in living patient-derived CML cells (Mizutani *et al.*, 2010; Horiguchi *et al.*, 2017). While monitoring kinase activity, we noticed that BCR-ABL is localized in liquid-like granules. Here, we identified that such BCR-ABL-positive granules are stress granules (SGs) and that their formation requires ABL kinase activity and the N-terminal region of BCR. Interestingly, its granular localization is necessary for the oncogenic function of BCR-ABL, demonstrating that a novel functional domain of BCR-ABL promotes the pathogenesis of CML.

## MATERIALS AND METHODS

### Reagents and antibodies

Imatinib was a kind gift from Novartis Pharma (Basel, Switzerland). Hoechst 33342 was obtained from Thermo Fisher Scientific (Carlsbad, CA, USA). Cycloheximide (used at a final concentration of 10 µg/ml) and thapsigargin (1 µM) were purchased from FUJIFILM Wako Pure Chemical Corporation (Osaka, Japan) and Sigma-Aldrich (St. Louis, MO, USA), respectively. An anti-c-Abl monoclonal antibody (sc-23) was obtained from Santa Cruz Biotechnology (Dallas, TX, USA). Antibodies against Hsp90α (ab2928) and Dcp1a (ab47811) were purchased from Abcam (Cambridge, UK). Alexa Fluor 488-, 594-, and 647-conjugated goat anti-mouse IgG antibodies and an Alexa Fluor 647-conjugated goat anti-rabbit IgG antibody were obtained from Thermo Fisher Scientific.

### Cell culture and transfection

HeLa (CCL-2) and Cos-1 (CRL-1650) cells were obtained from the American Type Culture Collection (Manassas, VA, USA). K562 (RCB1897), Ku812 (RCB0495), TOM-1 (ACC 578), ALL/MIK (CVCL_W883), MY (RCB1701), WEHI-3 (RCB0035) and Ba/F3-CL1 (RCB4474) cells were purchased from RIKEN CELL BANK (Ibaraki, Japan). K562 cells were maintained in RPMI 1640 (Sigma-Aldrich) supplemented with 10% fetal bovine serum (FBS, Thermo Fisher Scientific). Ku812, TOM-1, and ALL/MIK were cultured in RPMI 1640 containing 10% FBS and 1% sodium pyruvate. MY cells were maintained in minimal essential medium (MEM, Sigma-Aldrich) supplemented with 10% FBS and 1% sodium pyruvate. HeLa, Cos-1, and murine interleukin 3 (IL3)-producing WEHI-3 cells were cultured in Dulbecco’s modified Eagle medium (DMEM, Sigma-Aldrich) supplemented with 10% FBS. Ba/F3-CL1 cells were maintained in RPMI 1640 containing 10% FBS and IL-3-containing WEHI-3-conditioned media. The conditioned medium was recovered from near-confluent WEHI-3 cells, centrifuged for 5 min at 390 × *g*, and cleared by filtration with a 0.22 µm-diameter pore. All cells were maintained under a 5% CO_2_ humidified atmosphere at 37°C.

The expression vectors were transfected into HeLa, Cos-1, K562, Ku812, and TOM-1 cells with “Max” polyethylenimine (Polysciences, Warrington, PA, USA) according to the manufacturer’s recommendations. Gene transfer into K562, Ku812, TOM-1, and Ba/F3 cells was performed via nucleofection with the use of solution V and programs T-016 (for K562 cells), C-005 (for Ku812 and TOM-1 cells), and X-001 (for Ba/F3 cells) according to the manufacturer’s recommendations (Lonza, Basel, Switzerland).

### Plasmids

The expression vectors for BCR-ABL, Pickles 2.34, were described previously (Mizutani *et al.*, 2010; Horiguchi *et al.*, 2017). cDNA for BCR-ABL truncation mutants was generated by PCR with the following primers: BCR_F and BCR_fr1_R, BCR_fr2_F and BCR_fr2_R, BCR_fr3_F and BCR_fr3_R, BCR_fr4_F and BCR_fr4_R, BCR_fr5_F and BCR_fr5_R, BCR-ABL-J_F and BCR-ABL-J_R, ABL_fr1_F and ABL_fr1_R, ABL_fr2_F and ABL_fr2_R, ABL_fr3_F and ABL_fr3_R, ABL_fr4_F and ABL_fr4_R, and ABL_fr5_F and ABL_R. cDNA for BCR-ABL-ΔN was also obtained by PCR with BCR_fr3_F and ABL_R. The resulting PCR products were cleaved by *Xho*I and *Not*I and were then subcloned into the *Xho*I/*Not*I sites of the pFX-H2B-Venus (Kashiwagi *et al.*, submitted for publication).

cDNAs for FYVE, CD63, LAMP1, and TMEM192 were amplified by PCR with the following primers: FYVE_F and FYVE_R, CD63_F and CD63_R, LAMP1_F and LAMP1_R and TMEM192_F and TMEM192_R. These sequences were then subcloned into the *XhoI/NotI* sites of the pFX-mCherry vector or the pFX-TFP650 vector (Kashiwagi *et al.*, submitted for publication). mCherry-Sec61 β (# 49155) was obtained from Dr. Gia Voeltz via Addgene (Watertown, MA, USA). Expression vectors for other organelle markers will be described elsewhere (Kashiwagi *et al.*, submitted for publication). The primers used in this study are listed in Table S1.

### Microscopic setup

Cells were imaged with IX-83 and IX-81 inverted microscopes (Olympus, Tokyo, Japan) equipped with a BioPoint MAC 6000 filter and shutter control unit (Ludl Electronic Products, Hawthorne, NY, USA), an automated XY-stage (Chuo Precision Industrial, Tokyo, Japan), and a SOLA Light Engine (Lumencor, Beaverton, OR, USA) as an illumination source. UPlanSApo 60×/1.35 oil objective lenses were used. The following excitation and emission filters were used in this study: FF01-387/11-25 and FF02-447/60-25 (Semrock, Rochester, NY, USA) for Hoechst 33342; FF02-438/24 and FF01-483/32 (Semrock) for cyan fluorescent protein (CFP) and its derivatives; BP470-490 and BP510-550 (Olympus) for green fluorescent protein (GFP) derivatives; FF01-500/24-25 and FF01-542/27 (Semrock) for yellow fluorescent protein (YFP) and its derivatives; BP520-550 and BA580IF (Olympus) for red fluorescent protein (RFP) derivatives; and FF02-438/24 and FF01-542/27 (Semrock) for FRET. Confocal images were acquired with an sDISK spinning disk unit (Oxford Instruments, Abingdon, UK) and a Rolera EM-C^2^ electron multiplying cooled charge-coupled device camera (QImaging, Surrey, BC, Canada), whereas the epifluorescence images were acquired with a Cool SNAP MYO cooled charge-coupled device camera (Photometrics, Tucson, AZ, USA). MetaMorph software (Molecular Devices, San Jose, CA, USA) was used for the control of the microscope and the peripheral equipment. For live-cell imaging, the atmosphere was maintained at 37°C with a Chamlide incubator system (Live Cell Instrument, Seoul, Korea) for both microscopes.

### Live cell imaging

Cells expressing either fluorescent protein-tagged or epitope-tagged proteins were plated on 35-mm-diameter glass-bottom dishes (AGC Techno Glass, Shizuoka, Japan) coated with poly-L-lysine-(for hematopoietic cells) or collagen-coated (for nonhematopoietic cells). During the observation, cells were maintained in phenol red-free RPMI 1640 (for hematopoietic cells, Thermo Fisher Scientific) or phenol red-free DMEM/F12 (Thermo Fisher Scientific).

For imaging of FRET, cells were subjected to time-lapse, dual-emission microscopy with an interval of every 30 sec. Beginning at 10 min, cells were treated with 2 µM imatinib. To represent FRET efficacy, the FRET/CFP emission ratio was calculated as the quotient of background-subtracted FRET and CFP images and normalized to that at time 0. In Fig. 3, FRET/CFP ratio images are shown in intensity-modulated display mode, in which eight colors from red to blue represent the normalized emission ratios, with the intensity of each color indicating the mean intensity of FRET.

### Immunofluorescence

Cells were fixed with 3% paraformaldehyde for 15 min at room temperature. The cells were permeabilized with 0.1% Triton X-100 in PBS for 4 min at room temperature and then incubated with 1% bovine serum albumin and Hoechst 33342 (1 µg/ml). The cells were further incubated with primary antibodies (c-Abl, 1:1,000; Dcp1a, 1:1,000; Hsp90α, 1:200) overnight at 4ºC, after which immune complexes were detected by incubation for 1 h at room temperature in the dark with Alexa Fluor 488-, Alexa Fluor 594-, or Alexa Fluor 647-conjugated secondary antibodies (1:300). Images were acquired with an FV10i confocal microscope (Olympus).

### Quantification of BCR-ABL-GFP-positive granules

HeLa cells expressing BCR-ABL-GFP were either untreated or treated with 10 µg/ml cycloheximide for 2 h. The cells were then fixed, incubated with Hoechst 33342 and subjected to scanning confocal microscopy. Z-stack images were obtained from top to bottom of the cell with a step of 1 µm and then projected by maximum intensity. The number of granules was automatically counted from the projected images by the MetaMorph plugin “Granularity” within the area that was used for the determination according to the phase contrast images.

### Proliferation assay

Forty-eight hours after nucleofection, Ba/F3 cells were transferred into the media, including WEHI-conditioned media and 2 µg/ml puromycin, selected for 7-8 days. Stable cell lines were washed three times with prewarmed phosphate buffered saline (PBS), and 10^6^ live cells were plated in media devoid of IL-3. Viable cell counts were performed daily for 5 days with the trypan blue exclusion test. If necessary, cells were diluted appropriately to <1.0 × 10^6^ /ml.

### Statistical analyses

The quantitative data are presented as the mean ± standard error of the mean (s.e.m.) of at least three independent experiments (unless indicated otherwise) and were compared by one-way analysis of variance (ANOVA) followed by Student’s *t*-test (parametric test between two conditions). Time series data sets were compared by multivariate analysis of variance (MANOVA) with Bonferroni correction (among multiple conditions). No statistical methods were used to predetermine the sample size. The studies were performed unblinded.

## RESULTS

### BCR-ABL is localized to cytoplasmic granules

To investigate the subcellular localization of BCR-ABL in Ph^1^-positive cell lines, K562, Ku812, PALL-2, ALL/MIK, MY, and TOM-1 cells were fixed and subjected to immunofluorescence with the use of an anti-ABL antibody. We found that endogenous BCR-ABL was localized in the cytoplasm and accumulated in cytoplasmic granules in all cell lines (Fig. 1A), consistent with previous reports (Skourides *et al.*, 1999; Patel *et al.*, 2008).

**Fig. 1:**
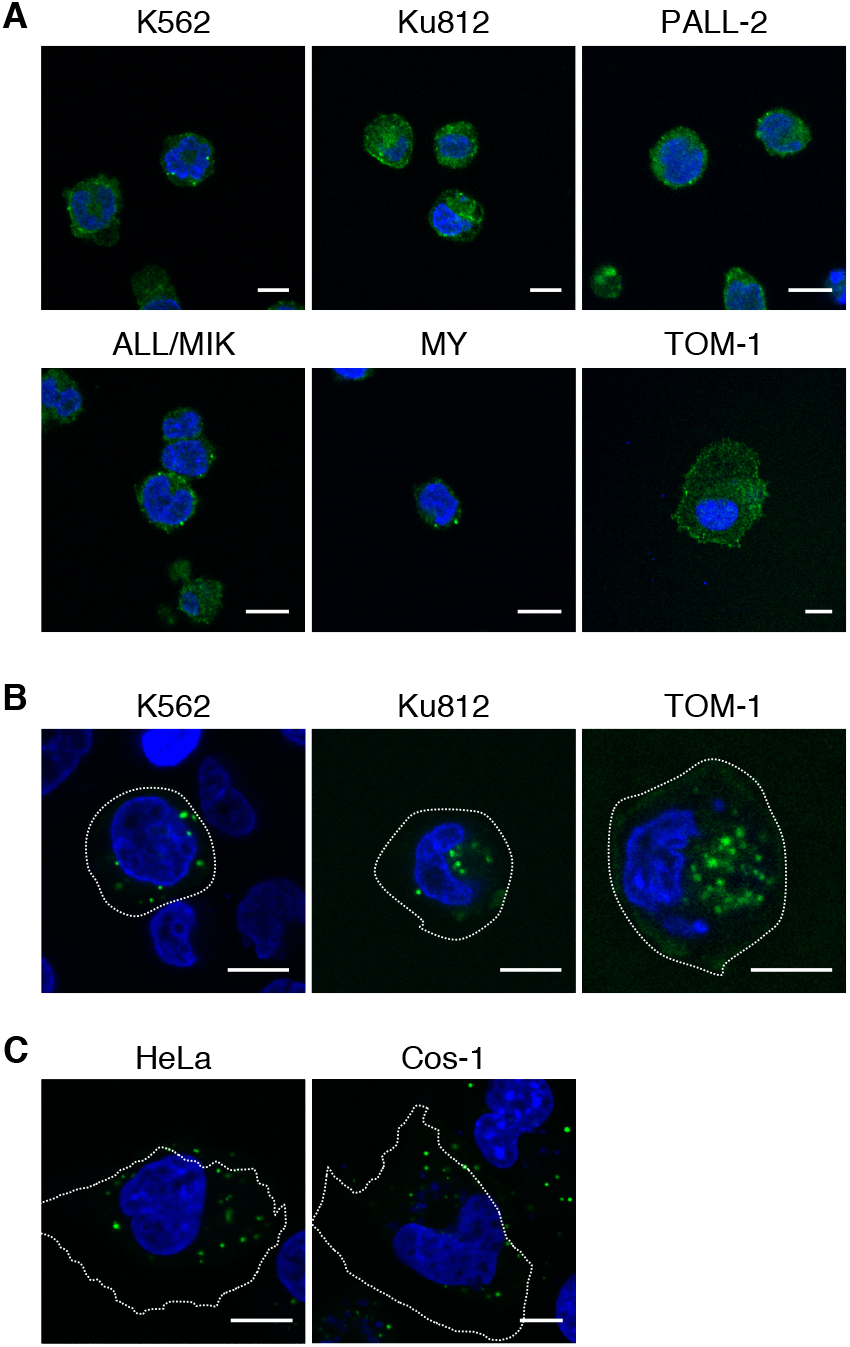
BCR-ABL is localized to granules. (A) Localization of endogenous BCR-ABL was analyzed by immunofluorescence using an anti-c-Abl antibody (green) in the cell lines indicated at the top. Nuclei were counterstained with Hoechst 33342 (blue). Representative confocal images are shown. Bar, 10 µm. (B, C) Expression vectors for BCR-ABL-GFP were transfected into the hematopoietic cell lines (B) or adherent cell lines (C) indicated at the top. The cells were observed with confocal microscopy. Representative images are shown. Outlines of cells are indicated by dashed lines. Bars, 10 µm.

Exogenously expressed BCR-ABL-GFP also formed clear cytoplasmic granules not only in the Ph^1^-positive cells (Fig. 1B) but also in nonhematopoietic cells (Fig. 1C). These results indicated that the formation of granules is due to the intrinsic properties of BCR-ABL but not to cell context-dependent conditions. It has been reported that BCR-ABL-positive granules were not colocalized with endosomal markers, EEA1, Rab7, transferrin, LysoTracker, or caveolin (Skourides *et al.*, 1999), encouraging us to examine in detail the localization of BCR-ABL with the use of a variety of organelle markers (Kashiwagi *et al.*, submitted for publication). Nevertheless, the BCR-ABL-GFP-positive granules were not colocalized with any markers for membrane-bound organelles, which include those for endosomes (early, late, and recycling endosomes), multivesicular bodies, lysosomes, endoplasmic reticulum, and mitochondria (Fig. S1).

### BCR-ABL accumulates in SGs

During live cell imaging of BCR-ABL-GFP, we noticed that the BCR-ABL-positive granules exhibited liquid-like behavior, i.e., fusion and separation (Fig. S2). It was reported that the heat shock protein Hsp90, one of the components of cytoplasmic RNA granules, including P-bodies (PBs) and SGs, interacts with BCR-ABL (An *et al.*, 2000; Matsumoto *et al.*, 2011; Pare *et al.*, 2009). Indeed, treatment with the translation inhibitor cycloheximide, which prevents RNA granule assembly through retention of messenger ribonucleoprotein polysomes (Mollet *et al.*, 2008), decreased the number of BCR-ABL-positive granules (Fig. 2A), confirming that BCR-ABL forms RNA granules. To further determine whether such RNA granules are PBs or SGs, the cells expressing BCR-ABL-GFP were fixed and subjected to immunostaining with the use of antibodies against HSP90α and the mRNA-decapping enzyme Dcp1a, a component of PBs. Endogenous Hsp90α accumulated in granular structures with BCR-ABL-GFP in HeLa cells, whereas it was localized diffusely in the cytoplasm in control, GFP-expressing cells (Fig. 2B). In addition, the colocalization of BCR-ABL-GFP and Hsp90α at the granular structures was also confirmed in K562 cells (Fig. S3A). In contrast, Dcp1a formed granular structures even in the absence of BCR-ABL, but such granules were not colocalized with BCR-ABL-GFP-positive granules (Fig. S3B). Moreover, thapsigargin treatment, which induces ER stress and subsequent SG formation (Thomas *et al.*, 2009), increased the number of BCR-ABL-GFP-positive granules (Fig. S3C). These results together suggest that BCR-ABL facilitates the formation of SGs and accumulates therein.

**Fig. 2:**
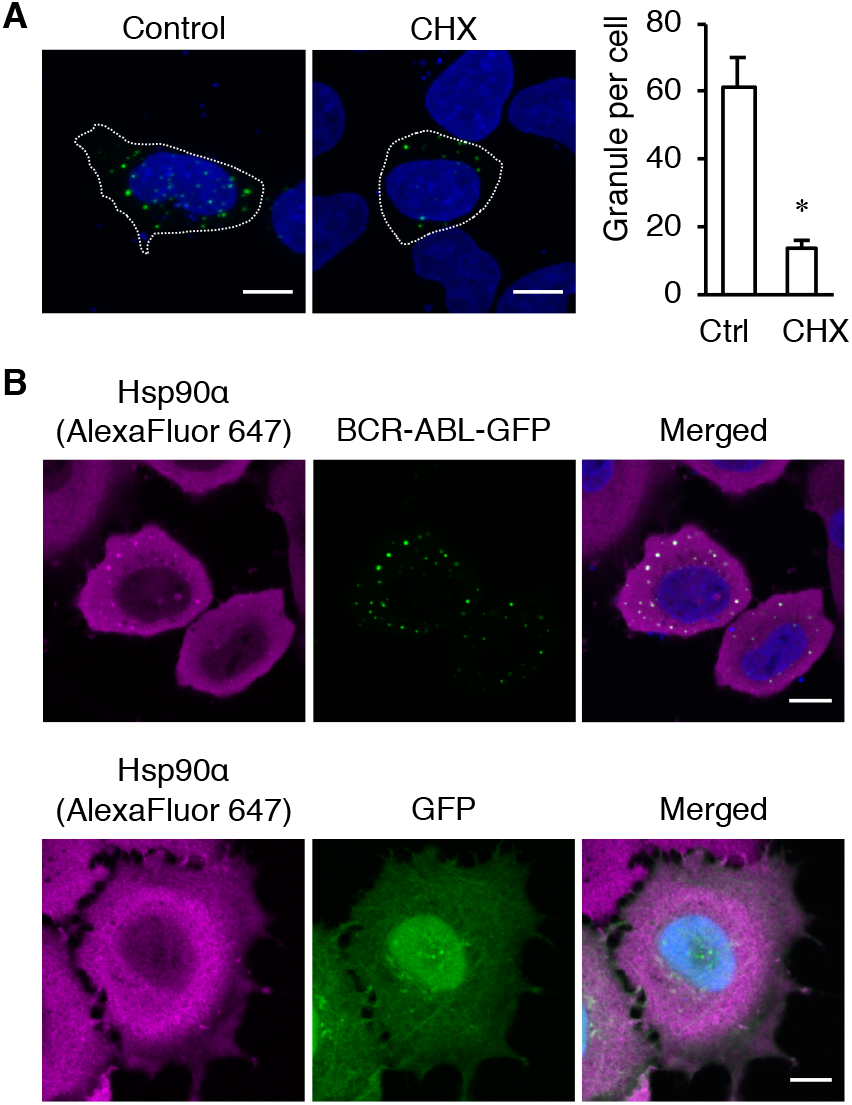
BCR-ABL is localized to RNA granules. (A) HeLa cells expressing BCR-ABL-GFP were either treated with cycloheximide (CHX) for 2 h or left untreated (control, Ctrl) and then observed with confocal microscopy. The maximum intensity projection of Z-stacks is shown. Nuclei were counterstained with Hoechst 33342 (blue). Outlines of cells are indicated by dashed lines. Bar, 10 µm. The number of cytoplasmic granules per cell was quantified as described in the Materials and Methods and plotted in the right graph. Data are means ± s.e.m. from three independent experiments. *, *p* < 0.0001 versus control as calculated by Student’s *t*-test. (B) K562 cells expressing GFP or BCR-ABL-GFP (green) were subjected to immunofluorescence with an anti-Hsp90α antibody (magenta). Nuclei were counterstained with Hoechst 33342 (blue). The cells were observed with confocal microscopy. Representative images are shown. Bars, 10 µm.

### Kinase activity of BCR-ABL is required for granule formation

It was previously reported that the kinase-deficient mutation (K1176R) abolishes the vesicle-like structure of BCR-ABL (Skourides *et al.*, 1999). In addition, another oncogenic fusion protein, nucleophosmin-anaplastic lymphoma kinase (NPM-ALK), was reported to form mRNA-containing cytoplasmic granules in a manner dependent on the kinase activity of ALK (Fawal *et al.*, 2006, 2011). We therefore compared the subcellular distribution of BCR-ABL in the presence or absence of the BCR-ABL tyrosine kinase inhibitor imatinib (Druker *et al.*, 1996). In the Ph^1^-positive cells, the number of BCR-ABL-positive granules was reduced by imatinib treatment (Fig. 3A). To provide more direct evidence for the relationship between the dynamics of kinase activity and granular localization, we utilized FRET-based biosensor Pickles to monitor BCR-ABL kinase activity in living cells (Mizutani *et al.*, 2010; Horiguchi *et al.*, 2017). Time-lapse microscopic observation revealed that imatinib inhibited the kinase activity immediately, followed by the disappearance of BCR-ABL-positive granules (Fig. 3B, C), indicating that BCR-ABL kinase activity is required for granule formation.

**Fig. 3:**
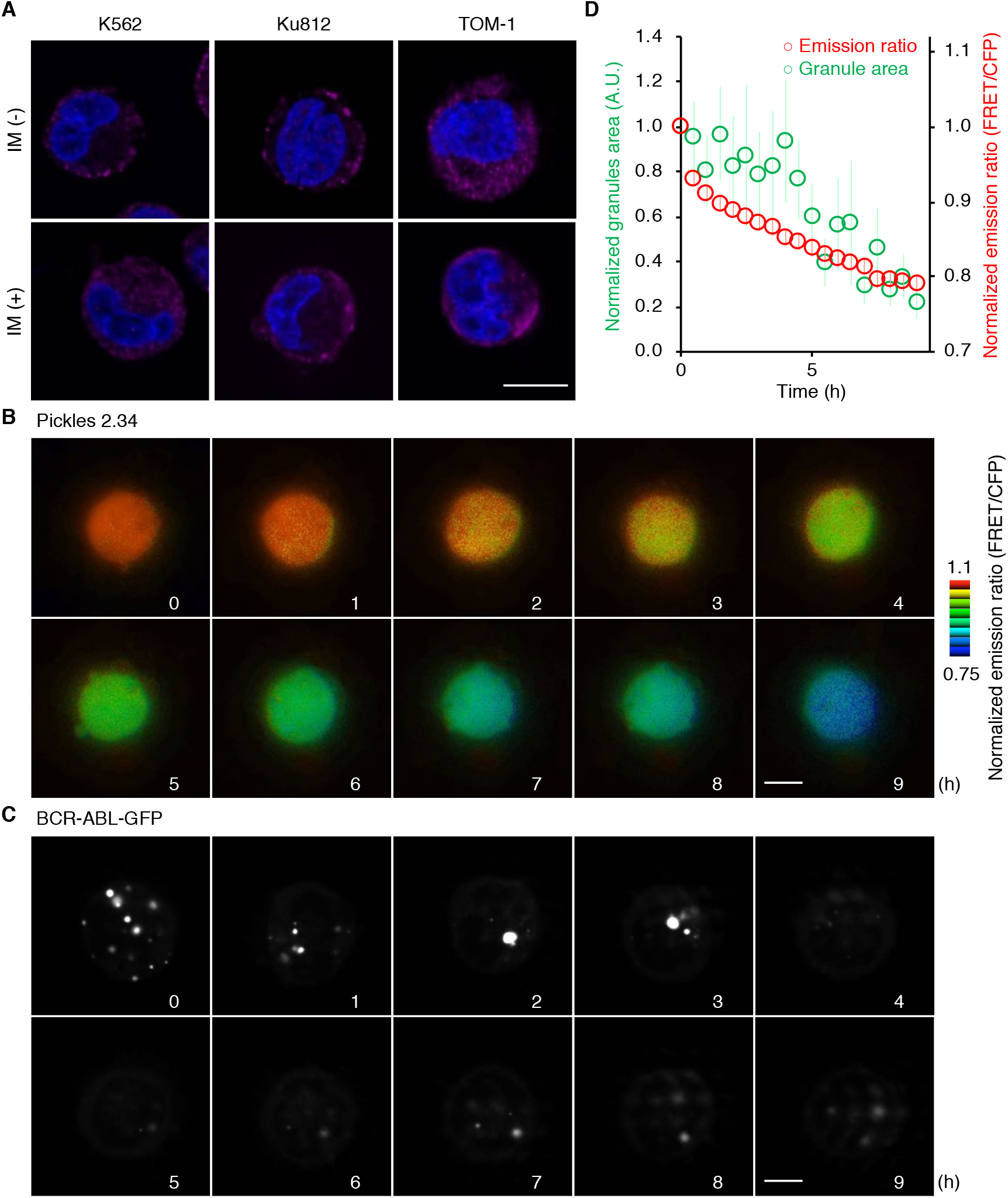
Tyrosine kinase activity is required for the granular localization of BCR-ABL. (A) The cell lines indicated at the top were either treated with 2 µM imatinib (IM) for 6 h or left untreated. The cells were then fixed and subjected to immunofluorescence with an anti-c-Abl antibody (magenta). Nuclei were counterstained with Hoechst 33342 (blue). (B–D) K562 cells expressing Pickles 2.34 (B) or BCR-ABL-GFP (C) were subjected to time-lapse microscopy. At time 0, the cells were exposed to IM. The emission ratio and the intensity of ECFP were used to generate the reconstituted images in the intensity-modulated display (IMD) mode. Representative images are shown over time (B). The time course of the emission ratio (red) and the number of granules (green) in the presence of IM were normalized and plotted over time (D). Data are means ± s.e.m. from four independent experiments.

### The N-terminal domain of BCR-ABL is necessary for its granular localization

To determine the domain responsible for granule formation, we next constructed a series of truncation mutants of BCR-ABL (Fig. 4A). Among the mutants, only the N-terminal fragment of BCR (BCR-fr1) clearly showed subcellular granules. This granular localization of BCR-fr1 was observed in both Ph^1^-positive and nonhematopoietic cells (Fig. 4B).

**Fig. 4:**
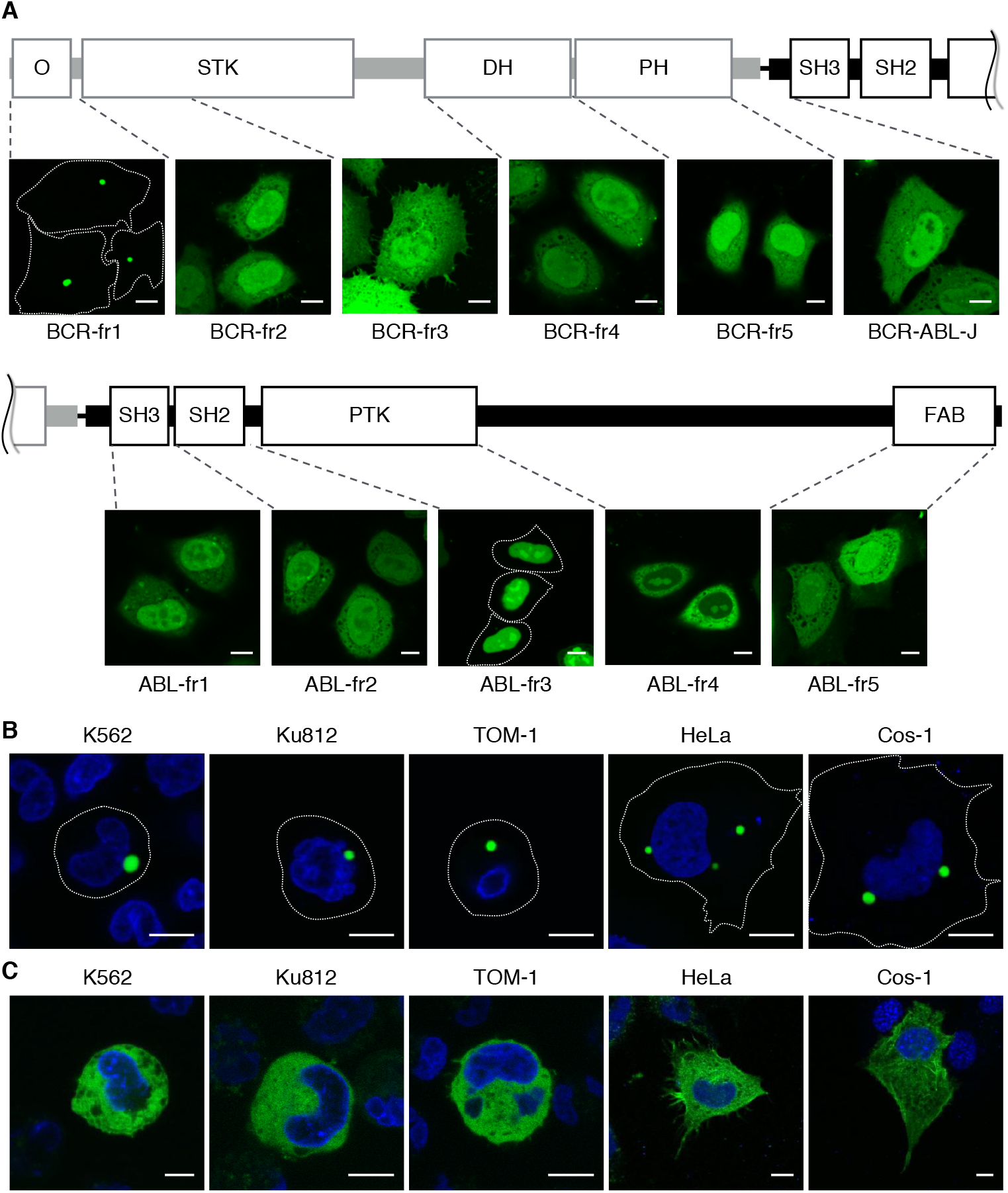
The N-terminal domain of BCR-ABL is responsible for its granular localization. (A) Schematic representation of the domain structure of BCR-ABL. Corresponding truncation mutants of BCR-ABL were fused with Venus and were expressed in HeLa cells. The cells were observed with confocal microscopy. Representative images are shown. (B, C) Expression vectors for Venus-BCR-fr1 (B, green) or BCR-ABL-ΔN-Venus (C, green) were transfected into the cell lines indicated at the top. The cells were observed with confocal microscopy. Representative images are shown. Bars, 10 µm.

Consistent with this finding, BCR-ABL lacking the N-terminal region of BCR (hereafter BCR-ABL-ΔN) was diffusely localized in the cytoplasm (Fig. 4C). These results suggested that the N-terminal region, namely, the CC domain, of BCR is involved in granule formation. Other mutants were localized diffusely in the cytoplasm and/or the nucleus (Fig. 4A).

***Cytosolic granule formation by BCR-ABL is essential for the proliferation of Ba/F3 cells*** It was reported that the N-terminal CC domain of BCR is required for the formation of homotetrameric BCR-ABL and its malignant transformation activity (Maru and Witte, 1991; Pendergast *et al.*, 1991; McWhirter *et al.*, 1993), although the mutants lacking this domain still possess tyrosine kinase activity (He et al., 2002). To clarify whether the oligomerization of BCR-ABL is necessary for granule formation, we utilized DsRed. DsRed forms an obligatory tetramer, which is required for the generation of red fluorescence (Sacchetti *et al.*, 2002). Therefore, tagging with DsRed promotes tetramerization of a covalently linked protein of interest. Moreover, the tetramerization can be confirmed by the emission of red fluorescence. Under these assumptions, whereas DsRed-tagged full-length BCR-ABL accumulated in cytosolic granules, DsRed-BCR-ABL-ΔN was diffusely localized in the cytoplasm (Fig. S4). Thus, the oligomerization of BCR-ABL is not sufficient for granule formation.

Finally, we examined whether granular localization is involved in the transformation activity of BCR-ABL. To this end, the lymphoid cell line Ba/F3 was utilized. Proliferation of this cell line requires IL-3, the depletion of which resulted in cell growth arrest (Mathey-Prevot et al., 1986, Fig. 5). Under this condition, whereas DsRed-BCR-ABL expression compensated for the growth defect, the expression of mCherry-BCR-ABL-ΔN, which is supposed to be in a monomer form, failed to do so (Fig. 5). Interestingly, DsRed-BCR-ABL-ΔN expression did not restore the defect (Fig. 5). Given that kinase activity and tetramerization, but not granule forming activity, of DsRed-BCR-ABL-ΔN was assumed to be conserved, these results indicated that the granular localization of BCR-ABL is necessary for its leukemogenic activity.

**Fig. 5:**
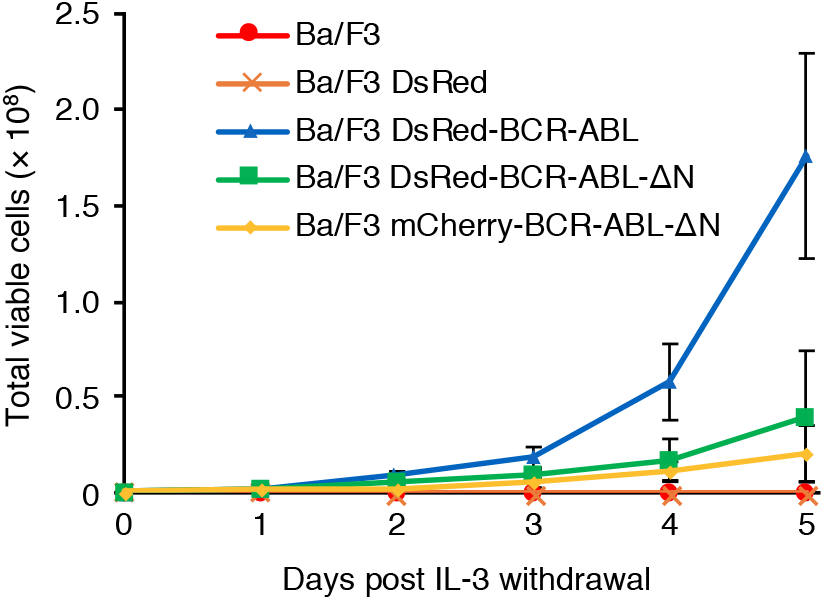
Tetramerization is not sufficient to increase the proliferation rate of IL-3-depleted Ba/F3 cells. Ba/F3 cells stably expressing the indicated proteins were cultured in medium containing IL-3 for three days before IL-3 depletion. At day 0, IL-3 was withdrawn from the media, and the number of cells was counted daily for 5 days. Note that the dead cells were excluded by trypan blue staining. Data shown are the mean ± s.e.m. from 3 independent experiments. *p*-values were calculated by MANOVA. *p* < 0.0001 for DsRed-BCR-ABL versus each other construct. Differences between other sets were not significant.

## DISCUSSION

In this study, we characterized cytosolic BCR-ABL-positive granules and demonstrated their significance for oncogenic activities. Such granules displayed liquid-like behavior and indeed colocalized with a marker for membrane-less compartments, SGs. Further investigations revealed that the formation of BCR-ABL-positive granules requires ABL kinase activity and the N-terminal region of BCR. Although it was previously reported that both endogenous and exogenous BCR-ABL form granular structures in CML cells and other cell lines, the subcellular localization of the granules has been controversial (Skourides *et al.*, 1999; Patel *et al.*, 2008). Indeed, no endosomal markers (e.g., markers for early or late endosomes) were colocalized with the BCR-ABL-positive granules. We have demonstrated that the BCR-ABL-positive granules are SGs; however, the molecular mechanism through which this fusion protein is localized to SGs has yet to be determined. In general, the biomolecular condensates, such as nucleoli, SGs, and PBs, are organized by liquid-liquid phase separation driven by interactions of multivalent molecules that harbor multiple elements for intra- or intermolecular interactions (Banani *et al.*, 2017). BCR-ABL is predicted to have the intrinsically disordered regions (IDRs), which provide the weak multivalency to promote phase separation (Banani *et al.*, 2017), in both BCR and ABL (1-417 and 1392-1916 of BCR-ABL) by the web-server-based disorder predictor IUPred (Maru, 2012). Therefore, it might be possible that these IDR domains are involved in granule formation by BCR-ABL. Another possibility might be the involvement of autophosphorylation of the Y177 residue in ABL kinase activity. Phosphorylated Y177 recruits Grb2, which participates in the phase separation of signaling molecules through multivalent interactions (Su *et al.*, 2016).

Similar to our findings on BCR-ABL, the chimeric protein NPM-ALK, a driver gene mutation for anaplastic large cell lymphoma, is also localized to cytoplasmic granules in a manner dependent on ALK kinase activity (Fawal *et al.*, 2006; Honorat *et al.*, 2006). These granules are identified as novel mRNA-containing cytoplasmic granules (ALK granules; AGs) and are distinct from PBs or SGs (Fawal *et al.*, 2011). Similar to our finding on BCR-ABL, AG formation requires ALK kinase activity (Fawal *et al.*, 2006), and NPM is involved in the phase separation through multivalent interactions (Banani *et al.*, 2017). However, it remains unclear whether and how formation of AGs contributes to ALK-mediated oncogenicity.

Non-membrane-bound compartments (e.g., nucleoli, SGs, and PBs) concentrate proteins and nucleic acids at discrete intracellular sites. This condensation could participate in fine-tuning of a variety of cellular processes (e.g., RNA metabolism, ribosome biogenesis, the DNA damage response, and signal transduction) through either the upregulation of reaction specificity and kinetics or inhibition by sequestration of relevant molecules (Banani *et al.*, 2017). For example, the DEAD-box helicase Ded1, a conserved component of SGs that promotes SG formation, represses translation of mRNAs accumulated in SGs (Hilliker *et al.*, 2011). In another example, Dishevelled 2 (Dvl2), a component of canonical and noncanonical Wnt pathways, accumulates in cellular granules (Schwarz-Romond *et al.*, 2007). The formation of Dvl2-positive granules might be required for activation of Wnt signaling because a mutant form of Dvl2 deficient in granule formation exhibits a dominant-negative effect on the Wnt response (Schwarz-Romond *et al.*, 2007). In the case of BCR-ABL, RNA metabolism and/or BCR-ABL activation in SGs is likely to be involved in tumorigenicity. In fact, it has been reported that the mRNA translation of the tumor suppressor breast cancer susceptibility gene 1 (BRCA1) is repressed via BCR-ABL-mediated SG formation in CML cells (Podszywalow-Bartnicka *et al.*, 2014). Additionally, a number of downstream signaling proteins (Grb2, p85 subunit of PI3K, SHC-transforming protein 1, E3 ubiquitin-protein ligase Cbl and CrkL) were found to be localized to the BCR-ABL-positive granules (Skourides *et al.*, 1999). Namely, given that the proportion of CrkL, a major substrate for leukemogenesis by BCR-ABL, is phosphorylated in the granules (Patel *et al.*, 2008), the concentration of the kinase and substrate (BCR-ABL and CrkL in this case) might facilitate excess phosphorylation of substrate and thereby promote leukemogenic activity.

In conclusion, we revealed that the phase separation of BCR-ABL contributes to its oncogenic function and that granule formation requires ABL kinase activity and the N-terminal region of BCR. Our findings suggest a novel mechanism for the oncogenicity of BCR-ABL: i.e., the N-terminal domain of BCR contributes oncogenicity by SG localization as well as by its oligomerization ability. Future studies may provide new therapeutic strategies based on the inhibition of BCR-ABL-positive granule formation.

## Author contributions

Conceptualization, Y.O.; investigation, S.K., Y.F., A.O.S., A.Y., M.F., H.S., T.K., T.T., and H.S.; writing, S.K. and Y.O.; funding acquisition, Y.F. and Y.O.; supervision, Y.O.

## ACKNOWLEDGEMENTS

We thank A. Miyawaki for the Venus cDNA, J. Groffen for the human CrkL cDNA, D. Baltimore for the BCR-ABL cDNA, Novartis Pharma for the IM, and A. Kikuchi for technical assistance. This work was supported in part by Grants-in-Aid from the Japan Society for the Promotion of Science (17H04016 and 19K22506 to Y.O.; 16H06227 to Y.F.) and from the Ministry of Education, Culture, Sports, Science and Technology (18H04850 and 19H05411 to Y.O.; 19H04823 to Y.F.), as well as by grants from the Canon Foundation, the Akiyama Life Science Foundation, the Nakatani Foundation, the Takeda Science Foundation, the Japan Research Foundation for Clinical Pharmacology, and the Tokyo Biochemical Research Foundation.

---

**Fig. S1:**
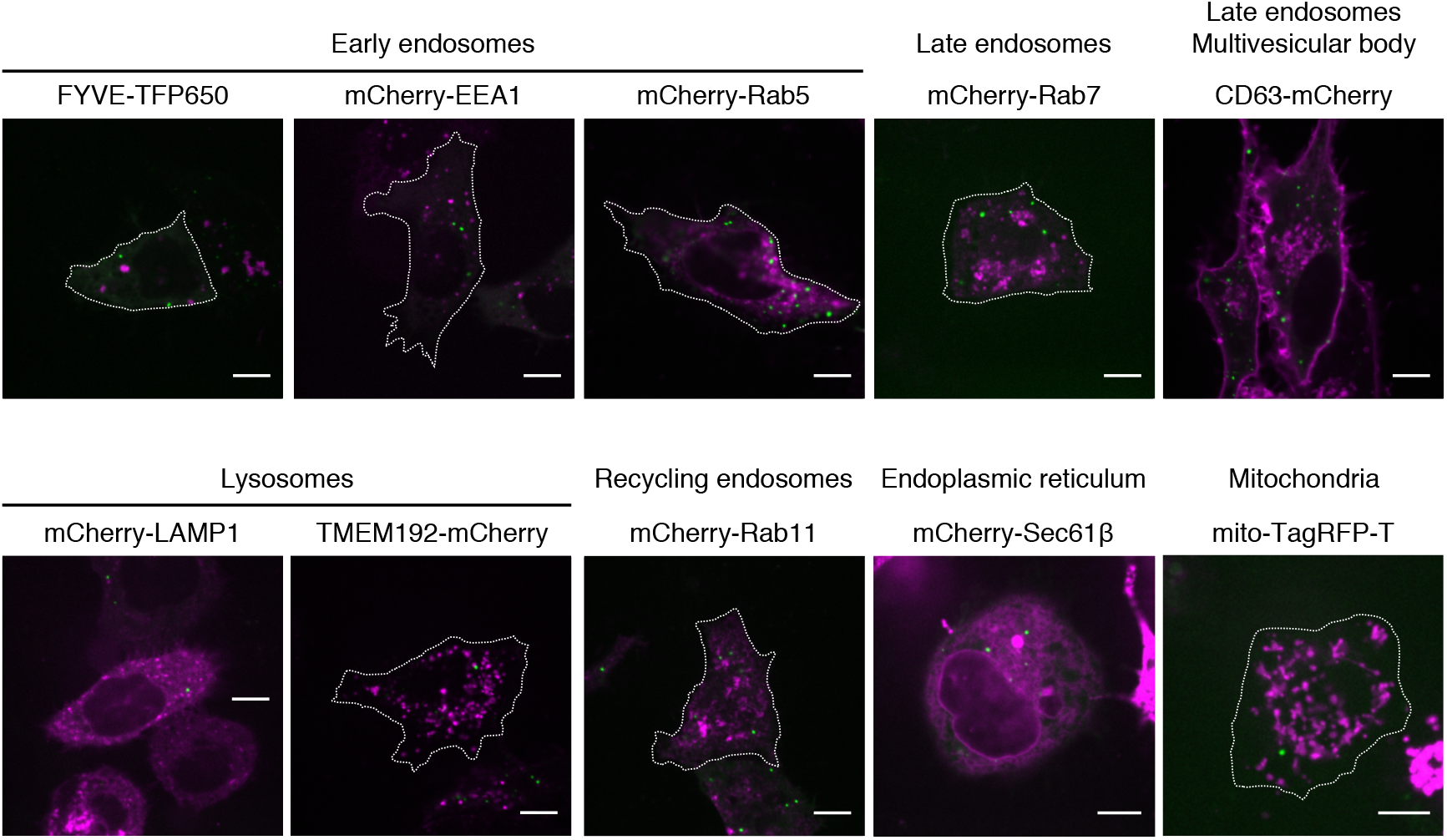
BCR-ABL is not localized to membrane-bound organelles. HeLa cells were transfected with expression vectors for BCR-ABL-GFP (green) and each organelle marker (magenta) indicated at the top. The cells were observed with confocal microscopy. Representative images are shown. Outlines of cells are indicated by dashed lines. Bars, 10 µm.

**Fig. S2:**
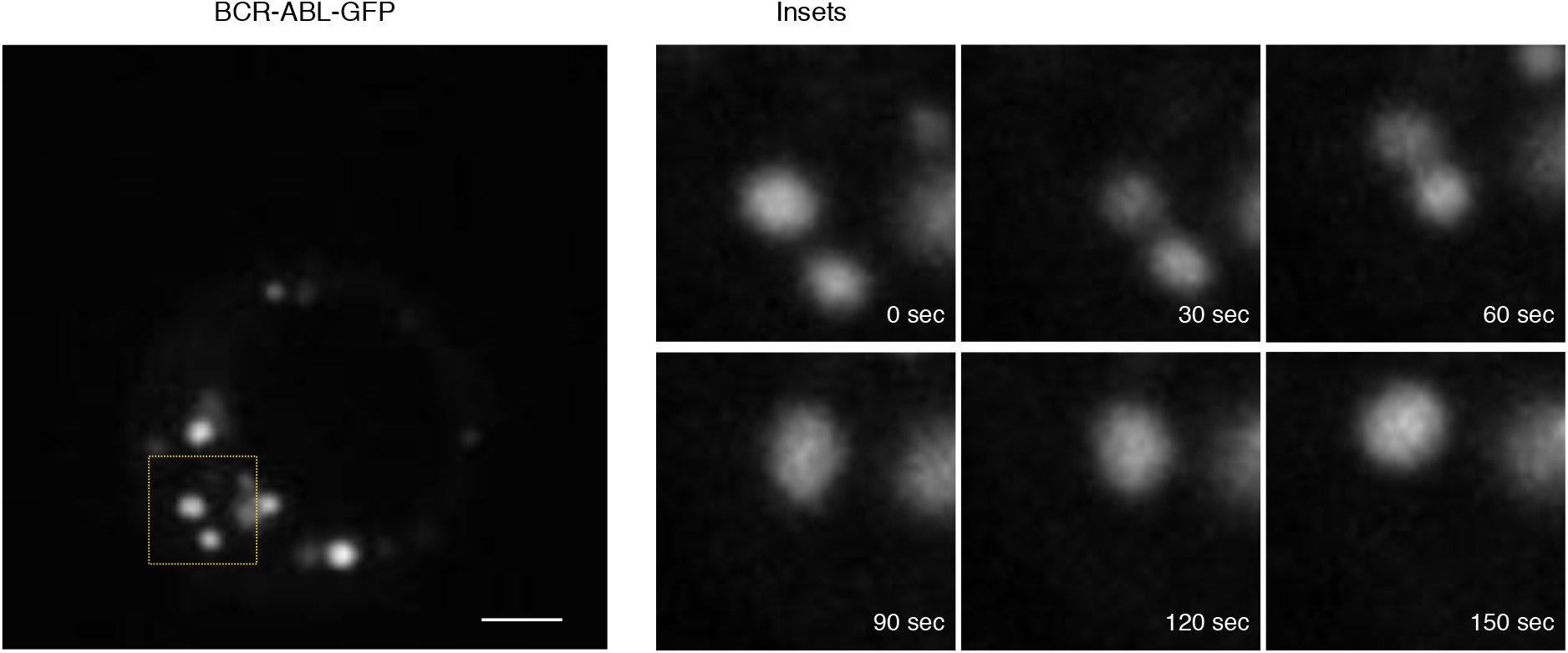
The BCR-ABL-positive granules exhibit liquid-like behavior. K562 cells were transfected with expression vectors for BCR-ABL-GFP and subjected to time-lapse confocal microscopy. The images were acquired every 30 sec. Right panels are higher magnification images of the inset indicated in the left entire-cell image. Representative images are shown. Bars, 10 µm.

**Fig. S3:**
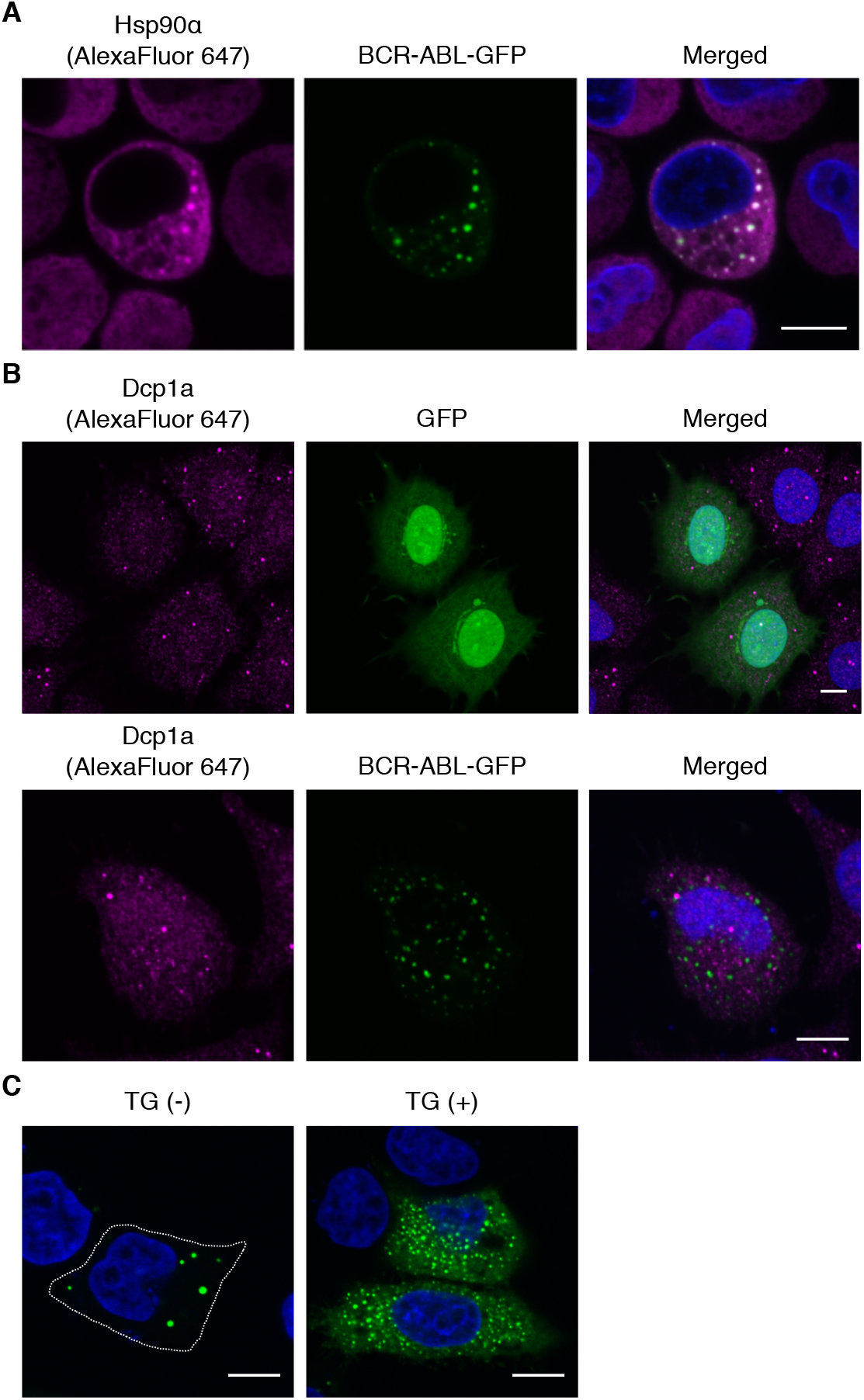
The BCR-ABL-positive granules are SGs. (A) K562 cells expressing BCR-ABL-GFP (green) were subjected to immunofluorescence with an anti-Hsp90α antibody (magenta). Nuclei were counterstained with Hoechst 33342 (blue). The cells were observed with confocal microscopy. Representative images are shown. Bars, 10 µm. (B) HeLa cells were transfected with BCR-ABL-GFP (green) for 24 h and subjected to immunofluorescence with an anti-Dcp1a antibody (magenta). Nuclei were counterstained with Hoechst 33342 (blue). The cells were observed with confocal microscopy. Representative images are shown. Bars, 10 µm. (C) HeLa cells expressing BCR-ABL-GFP were either treated with thapsigargin for 1 h [TG (+)] or left untreated [TG (-)] and then fixed and observed with confocal microscopy. Representative images are shown. Outlines of cells are indicated by dashed lines. Bars, 10 µm.

**Fig. S4:**
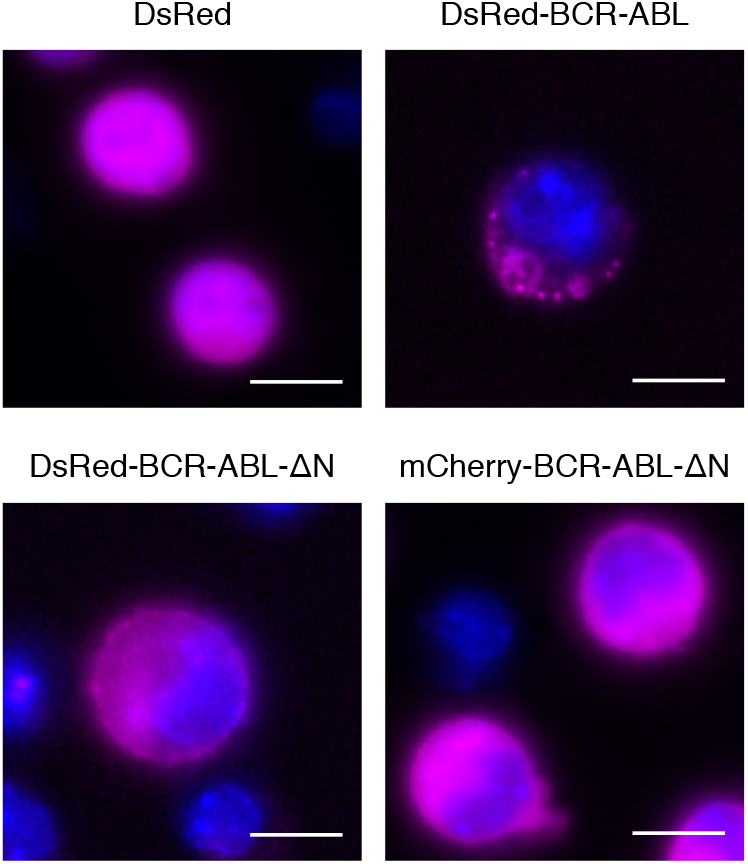
Tetramerization of BCR-ABL is not involved in its granular localization. Ba/F3 cells stably expressing the indicated proteins (magenta) were observed with epifluorescence microscopy. Nuclei were counterstained with Hoechst 33342 (blue). Representative images are shown. Bars, 10 µm.

**Supplementary Table 1.**
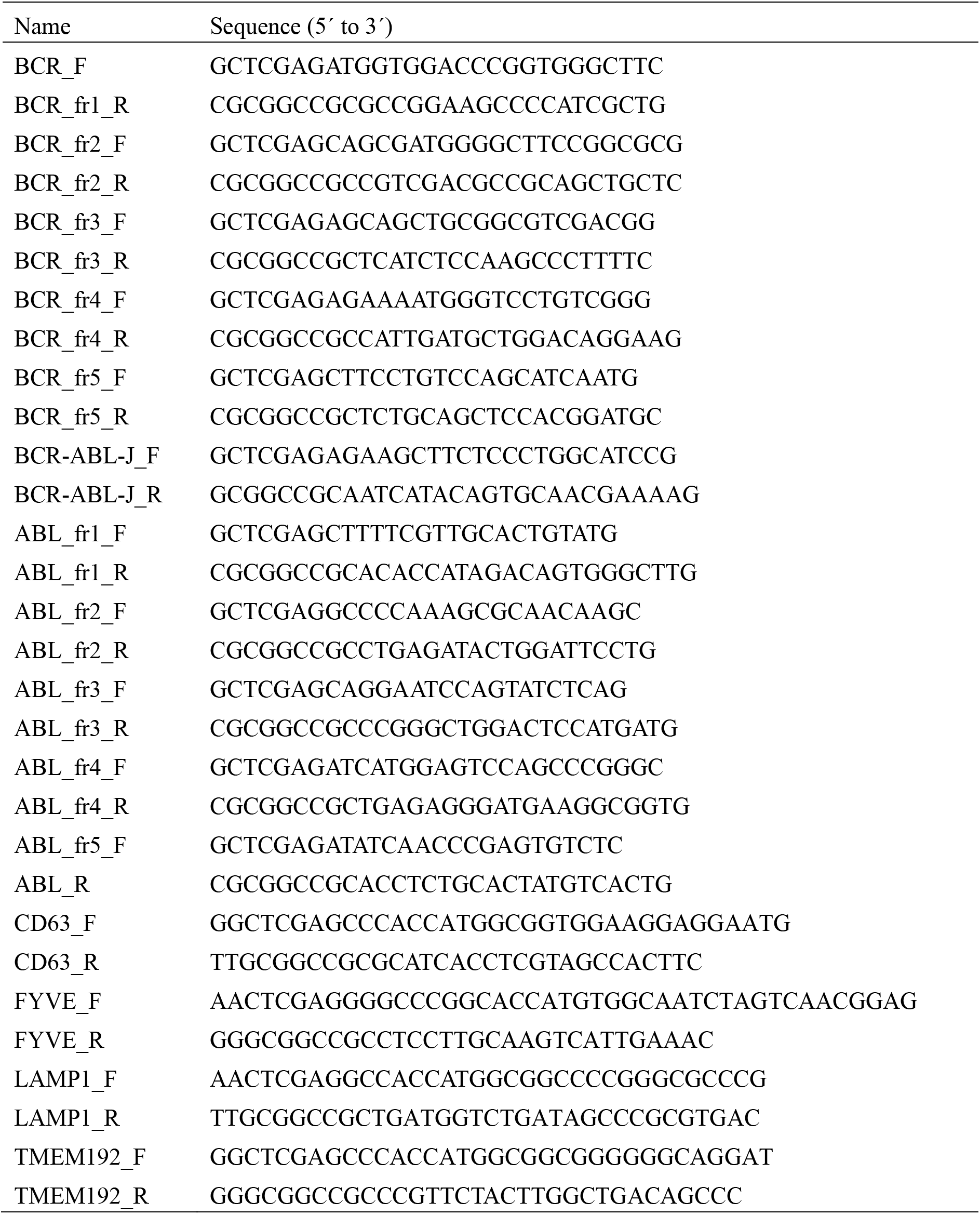
Primers used in this study.

